# Explainable deep graph learning accurately modeling the peptide secondary structure prediction

**DOI:** 10.1101/2022.06.09.495580

**Authors:** Yi Jiang, Ruheng Wang, Jiuxin Feng, Junru Jin, Sirui Liang, Zhongshen Li, Yingying Yu, Anjun Ma, Ran Su, Quan Zou, Qin Ma, Leyi Wei

## Abstract

Accurately predicting peptide secondary structures remains a challenging task due to the lack of discriminative information in short peptides. In this study, we propose PHAT, a deep graph learning framework for the prediction of peptide secondary structures. The framework includes a novel interpretable deep hypergraph multi-head attention network that uses residue-based reasoning for structure prediction. Our algorithm can incorporate sequential semantic information from large-scale biological corpus and structural semantic information from multi-scale structural segmentation, leading to better accuracy and interpretability even with extremely short peptides. Our interpretable models are able to highlight the reasoning of structural feature representations and the classification of secondary substructures. We further demonstrate the importance of secondary structures in peptide tertiary structure reconstruction and downstream functional analysis, highlighting the versatility of our models. To facilitate the use of our model, we establish an online server which is accessible via http://inner.wei-group.net/PHAT/. We expect our work to assist in the design of functional peptides and contribute to the advancement of structural biology research.

## Introduction

Peptides have recently emerged as potential therapeutic molecules against various diseases, and have garnered increasing attention due to their many advantages, including high specificity, high penetration, low production cost, and ease of manufacturing and modification [1]. Various disease-specific functional peptides have entered the global market, including antiviral peptides (AVPs), antimicrobial peptides (AMPs), and anticancer peptides (ACPs) [2–4]. Specifically, a family of peptides known as cell-penetrating peptides (CPPs) has shown enormous success in the cellular uptake of therapeutic molecules [5]. Currently, over 40 cyclic peptide drugs are in clinical use, and approximately one new cyclic peptide drug is approved for clinical use each year on average [6]. Furthermore, predicting the secondary structure of bioactive peptides can provide key insights into the functional mechanisms of peptides and could serve as a basis for designing peptides with desired functions [1]. Predicting the secondary structure of peptides is an intermediate step in predicting three-dimensional (3D) or tertiary structures, all of which are essential determinants of peptide bioactivity [7]. Therefore, reliable and accurate computational methods for predicting the secondary structures of peptides are urgently needed.

Many efforts have been made to predict the secondary structure of proteins through computational approaches, most of which are based on machine learning algorithms. For instance, Heffernan *et al*. developed a multi-task deep learning model [8] in which a long- and short-term memory bidirectional regression neural network (LSTM-BRNNS) was constructed to capture both short-term and long-term residue interaction relationships [9]. Li *et al*.developed the diffusion convolutional recurrent neural network (DCRNN), a hybrid neural network that alleviates the local features derived from convolutional neural networks (CNNs) and the global features captured from stacked bi-directional gated recurrent units (BIGRU) to predict the secondary structures of proteins [10]. Similarly, Busia *et al*. integrated CNN and residual connections to predict the secondary structures of peptides and achieved good performance, demonstrating the importance of the primary protein sequence information in secondary structure prediction [11]. In addition to the above methods, there are many other protein secondary structure predictors, such as DeepCNF, JPRED, PROTEUS2, RaptorX, and MUfold-SSW, among others [12–17]. However, these methods are designed for the prediction of protein structures and are not applicable for secondary structure prediction due to the inherent structural differences between peptides and proteins. For example, evolutionary information is frequently integrated and used for model training in the prediction of protein secondary structures, and potential biases might be introduced when designing peptide secondary structure models due to the short length of peptides. Additionally, previous studies have demonstrated that even for identical segments of residues in proteins and peptides, their secondary structures might be quite different [1]. One possible reason is that proteins have more complex tertiary structures, which presumably leads to changes in secondary structures. Particularly, hydrophobic collapse is a major force responsible for a well-defined tertiary structure. However, this phenomenon is only applicable to proteins and not peptides [18]. Therefore, developing a peptide-specific secondary structure prediction method is urgently needed.

Singh *et al*. [1] proposed PEP2D, the first peptide-specific secondary structure predictor that trains a random forest (RF) model with peptide sequential and evolutionary data and achieves good performance. Recently, Cao *et al*. [19] designed PSSP-MVIRT (**P**eptide **S**econdary **S**tructure **P**rediction based on **M**ulti-**V**iew **I**nformation, **R**estriction and **T**ransfer learning) for the prediction of peptide secondary structures, employing CNNs and BIGRU to learn high-latent features and introducing transfer learning to overcome the lack of training data. In addition to the aforementioned methods, there are several other peptide structure prediction methods, such as PEP-FOLD [20]. However, existing methods have several limitations. Particularly, most of them rely heavily on feature engineering to design handcrafted features, the quality of which might greatly impact the predictive performance because the feature design is based on the researchers’ prior knowledge. Additionally, existing protein-specific secondary structure prediction methods focus on long-distance dependence of sequences with hundreds of residues rather than local fragments, whereas peptide-specific methods focus more on neighborhood information among residues, thus easily ignoring global information. Ultimately, although deep learning has been successfully used in secondary structure prediction, the current methods still follow a “black box” model and lack good interpretability. These shortcomings limit our ability to predict the relationships between peptide primary sequences and their secondary structures.

In this study, we propose an innovative deep learning model called PHAT to predict peptide secondary structures. Importantly, our proposed model incorporates several novel features: (*i*) we introduce a powerful pre-trained protein language model [21] to transfer semantic knowledge from large-scale proteins to peptides and learn high-latent and long-term features of peptide residues. (*ii*) Considering the local continuity and diversity of peptide secondary structures [22, 23], we propose a novel HyperGMA (**Hyper G**raph **M**ulti-head **At**tention network), in which we can encode peptide residues with multi-semantic secondary structural information while capturing contextual features from consecutive regions using multi-level attention mechanisms. Additionally, our constructed hypergraph effectively prevents over-smoothing, which is a common issue in conventional graph networks (*e.g*., GCN [24], GAT [25]). (*iii*) To reveal the predicting mechanisms of PHAT, the transition and emission matrices were visualized in conditional random fields (CRFs) that can automatically learn a set of biologically meaningful knowledge on secondary sub-structures. This overcomes the limitations of “black-box” approaches in deep learning-based models to some extent and provides good interpretability of our PHAT model. (*iv*) We also demonstrated that the structural predictions obtained from our model can assist in peptide-related downstream tasks, such as the prediction of peptide toxicity [26], T-cell receptor (TCR) interactions with MHC (major histocompatibility complex)-peptide complexes [27], and protein-peptide binding sites. (*v*) A case study demonstrated that our PHAT can also accurately predict distance map and contact map matrices, which can be further used for the reconstruction of peptide 3-D structures. Benchmarking results indicated that the proposed PHAT significantly outperforms the state-of-the-art methods in either 3-state or 8-state secondary structure prediction, demonstrating the superiority and robustness of our model. To facilitate the use of our method, we established a code-free, interactive, and non-programmatic web interface of PHAT at http://inner.wei-group.net/PHAT/, which can lessen the programming burden for biological and biomedical researchers.

## Materials and methods

### Datasets

To evaluate the effectiveness of our model, we used the same benchmark dataset commonly used as a “gold standard” dataset in several studies [19, 28]. The dataset contains 5,772 secondary structures of protein data with three structural states: Helix (H), Strand (E), and Coil (C). The dataset processing process is illustrated in **Figure 1A**. Specifically, the protein structures are derived from X-ray crystallography, and this process is executed with a resolution of at least 2.5 Å, with no chain breaks and less than five unknown amino acids. The sequence similarity in this dataset is reduced to 25% to ensure a fair performance evaluation. Additionally, there are some sequences containing the “X” symbol, representing unnatural residues in this dataset. Following the same data pre-processing in [19], we removed the unnatural sequences including the “X” symbol, and 4,542 protein and peptide sequences were retained. Afterward, among the remaining sequences, we selected the sequences with <100 residues lengths, finally yielding 1,285 peptide sequences as our three-structure-state dataset. The length of the sequences ranged from 30 to 99 residues. Moreover, previous studies have demonstrated that the secondary structures of protein and peptides can also be defined with eight states, including H (alpha-helix), G (3_10_helix), I (π-helix), E (extended beta-strand), B (isolated beta-strand), T (turns), S (bend), and others (C) [8, 29, 30]. To account for this scenario, we further constructed a new dataset of 1,060 peptide sequences, derived from the DSSP (Dictionary of Protein Secondary Structure) structure database [31].

**Figure 1.**
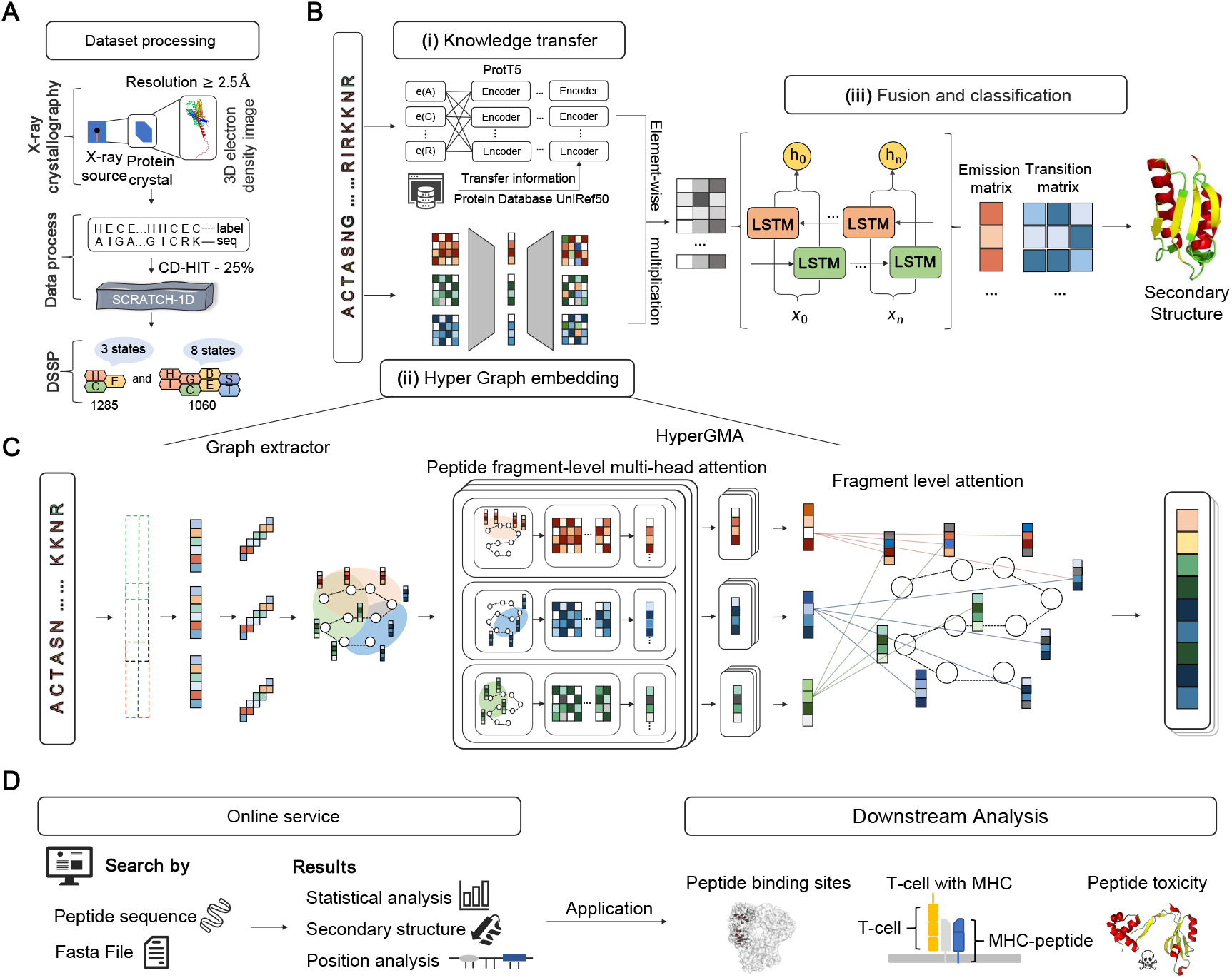
The workflow and framework of PHAT. (**A**) Dataset processing. We extracted the benchmark datasets from SCRATCH-1D, where the protein and peptide structures were derived with X-ray crystallography and operated with a resolution of at least 2.5 angstroms, for three-state and eight-state secondary structures. (**B**) Framework of PHAT. The framework consists of three modules: (*i*) Knowledge transfer module, (*ii*) Hyper Graph embedding module, and (*iii*) Fusion and classification module. In Knowledge transfer module, the original sequences are encoded by a pretrained protein model to gain the features of peptide residues. In Hyper Graph embedding module, the peptide sequences are constructed into hypergraph structures and embedded by HyperGMA. In Fusion and classification module, the outputs of Knowledge transfer module and the Hyper Graph embedding module are firstly fused through the element-wise multiplication and better integrated by the Bi-LSTM. Then the output of Bi-LSTM is inputted into the CRF layer, and as a result, the secondary structure of related residues can be predicted. (**C**) illustrates the details of Hyper Graph embedding module. In the part of graph extractor, peptide sequences are firstly sliced into fragments with specific length and constructed as hyperedges of the hypergraph structure. Then the hyperedges are cut into residue groups to be built as hypernodes in the hypergraph structure. Next, the hypergraph structure from graph extractor is inputted into HyperGMA to capture the multi-scale relationships in view of residue groups and peptide fragments by the multi-scale attention mechanism. (**D**) Online service. Our web server of PHAT is freely available to provide researchers with peptide details in three-state or eight-state secondary structures, statistical analysis, and position analysis. The predictions of our model can be applied in many downstream tasks as in Downstream Analysis.

### Training and testing datasets

To account for the characteristics of short peptide sequences and fairly evaluate the performance of the methods, the dataset was divided into two categories: >50 residue sequences and ≤50 residue sequences. The sequences with ≤50 residues consisting of 257 peptide sequences (with H of 5,294, E of 1,119, and C of 3,733) were used as the test set. The remaining 1,028 peptide sequences were used as the training dataset. For model training, we randomly selected 10% peptide sequences as our validation set from the training dataset to adjust the parameters of our model. Additionally, the training and testing datasets were labeled with the three-state secondary structures, with the sequence length of peptides ranging from 30 to 100. For the eight-structure-state dataset, we also collected 1,060 sequences to re-train and test our model. The details of the datasets are summarized in **Supplementary Table 1** and **Supplementary Table 2**.

### Architecture of the proposed PHAT model

The overall network architecture of the PHAT model is illustrated in **Figure 1B** with three main modules: (*i*) knowledge transfer module, (*ii*) hypergraph embedding module, and (*iii*) feature fusion and classification module. Specifically, our PHAT model only takes peptide sequences as input. In module (*i*), to address the scarcity of peptides, our model employs and fine-tuned and pre-trained large-scale protein language model called ProtT5 for the analysis of our peptide datasets. By doing so, we can transfer rich contextual information from large-scale protein sequences to our model and learn discriminative feature embeddings of peptide sequences. In module (*ii*), we propose a HyperGMA (**Hyper G**raph **M**ulti-head **A**ttention network) to learn local and global features. Specifically, given a peptide sequence, our model first exploits the graph extractor to divide the peptide sequence into fragments with particular lengths as hyperedges and residue groups as hypernodes. Then, by using the hyperedges and hypernodes, we construct the hypergraph structure and pass it to the HyperGMA to integrate the sequence information of different scales in the hypergraph structure. Our model can capture both local and global features at the residue group level and peptide fragment level through the multi-scale hypergraph attention mechanism. Afterward, in module (*iii*), we integrate the feature embeddings from the above two channels (knowledge transfer module and hypergraph embedding module) through an element-wise multiplication strategy. Furthermore, our model adopts Bi-LSTM (Bidirectional Long Short-Term Memory Networks) [32] to improve and optimize the feature representation ability and exploits CRFs to learn useful correlations among the sub-secondary structures. Finally, PHAT takes the resulting features from module (*iii*) as the input of a Viterbi algorithm and predicts the structural state to which each peptide residue belongs.

### Feature embedding from the pre-trained model ProtT5

Although there are some differences between proteins and peptides in terms of structure, they are similar in many aspects such as the transcription process and residue sequence composition. Therefore, we used the pre-trained model ProtT5 based on the t5-3b model [33], which was pre-trained using the UniRef50 database [34] (i.e., a database consisting of 45 million protein sequences), in a self-supervised manner to transfer semantic knowledge from proteins to peptides. Its weight was pre-trained with a BERT-like mask language model denoising objective using raw protein sequences without labeling. The model can fully learn the semantic information and generate different residue features belonging to multiple expressions in different context scenarios.

The original peptide sequences are fed into ProtT5, and the output vectors are extracted from many encoder blocks that are dependent on the self-attention mechanism. Each encoder block calculates the attention for each residue with all residues in the sequence, aiming to obtain the relevance and importance between every two residues. The calculation formula of the self-attention mechanism is as follows:

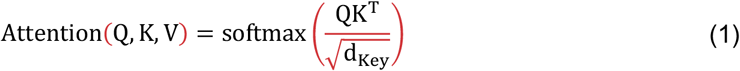

where Q, K, and V are the query vector, key vector, and value vector of the corresponding individual residues in the peptide sequence, respectively, and d_Key_ is the dimension of the input key vector.

### Hypergraph multi-head attention networks

Inspired by the previous studies for hypergraphs in natural language processing [35], we constructed a hypergraph structure by taking the peptide residue groups as nodes and the peptide fragments as edges. Based on this structure, we proposed a novel HyperGMA. **Figure 1C** shows the hypergraph construction process and HyperGMA architecture. (*i*) The peptide sequence was inputted into the graph extractor, which takes a particular length as the sliding window size and moves the sliding window to select the sequence fragments with cross residues. (*ii*) The sequence fragment is divided into smaller residue groups in a similar way as in step (*i*) but with a smaller sliding window size. The residue groups are regarded as hypernodes and the peptide fragments are taken as the hyperedges. (*iii*) The structure of the hypergraph is constructed using the hyperedges and hypernodes generated from steps (*i*) and (*ii*). (*iv*) Then, the hypergraph structure is inputted into HyperGMA to extract the graph embeddings of the peptide sequence.

The context of residues in a peptide sequence describes the language characteristics of local co-occurrence among residues, and its function in sequence representation learning has also been proved to be effective. In our model, we established two residues as a group, based on which we identified 400 types of groups. Moreover, a set of residue groups is regarded as a hyperedge, which is a sequence fragment with a specific length. This enables our model to simultaneously capture structural information both at the residue group level and peptide fragment level. Specifically, a hypergraph is defined as *G* = (*v, ε*), where *v* = {*v*_1_,*v*_2_,…,*v*_n_} represents a set of *n* nodes in the graph, and *ε* = {*e*_1_,*e*_2_,…,*e*_m_} represents a set of *m* hyperedges. Moreover, the model can connect two or more nodes for any hyperedge e_j_.

### Residue group-level multi-head attention

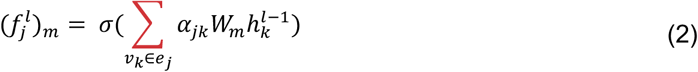

where *k* represents the index of the residue group (hypernode) in the fragment (hyperedge) *e_j_*, *j* indicates the index of the fragment in edge set *ε*, *v_k_* ∈ *e_j_* indicates that *v_k_* is contained in fragment e_j_, 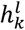 is the representation of residue group (hypernode) *v_k_* at layer *I, σ* is the activation function *LeakyReLU*, *W_m_* is the weight matrix trained in the m-head attention, and *m* represents the head number of multi heads. *α_jk_* is the attention coefficient of the residue group *v_k_* in the fragment e_j_, which can be computed by:

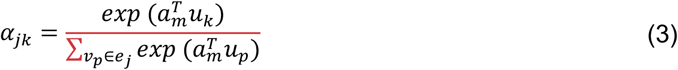

where 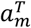 is a weight vector for measuring the importance of residue groups in the m-head attention, *v_p_* ∈ *e_j_* represents that residue group *v_p_* is contained in fragment *e_j_*, and *T* means “transpose.”*u_k_* represents *v_k_* on the hypergraph defined as:

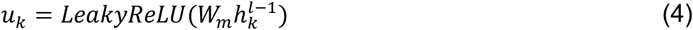

The expression 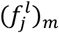 represents hyperedge *e_j_* from *m*-head attention at layer *l*. We constructed the multi-head attention mechanism, connected it, and compressed it to the desired dimension after the layer was fully connected. This structure is aimed to capture residue context information. The output 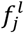 represents the connected representation of hyperedge e_j_ at layer *l*.

### Peptide fragment-level attention

With the representations of all peptide fragments (hyperedges) as 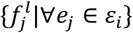 connecting to residue group *v_i_*, we introduce the fragment level attention mechanism to capture the structural information of peptide fragments with distance interval for learning the next-layer representation of residue group *v_i_*, which is expressed as follows:

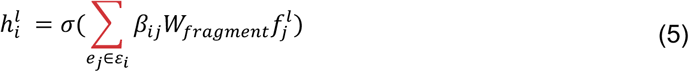

where 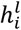 is the output representation of residue group (hypernode) *v_i_* (*v_i_* ∈ *v*) at layer *l, i* represents the index of the residue group (hypernode) in the node set *v*, all the hyperedges containing residue group *v_i_* are in *ε_i_*, and *W*_fragment_ is a weight matrix. *e_j_* is a fragment (hyperedge) divided at a fixed length from peptide sequence, and *ε_i_* is the set of fragments of the peptide.

*β_i,j_* shows the attention interaction of peptide fragment (hyperedge) *e_j_* on residue group (hypernode) *v_i_*. The computing process is described below:

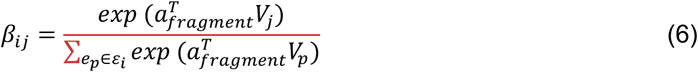

where 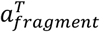 is a weight vector similar to 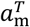 but for measuring the importance of peptide fragments

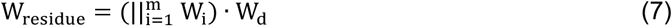

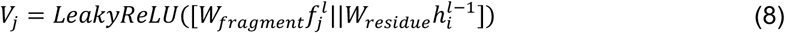

|| represents the concatenation operation, · is matrix multiplication, and *W_d_* is a trainable matrix for dimensional reduction.

### Bidirectional long short-term memory and conditional random field

The secondary structural information in peptide sequences is often related to the residues in the forward and backward peptide fragments. Therefore, we implemented Bi-LSTM (Bidirectional Long Short-Term Memory Networks) to extract information from two directions in the peptide sequence. Additionally, the previously learned features from ProtT5 and HyperGMA are fused in the form of element-wise multiplication, which may introduce redundant information. Therefore, we added a layer of Bi-LSTM to better integrate them and provide a sequence-level view for the CRF layer. Bi-LSTM is a deep-learning architecture with two LSTM layers in different directions, which can capture the dependence of long-distance residues, and selectively learn and forget information with corresponding importance through training [36]. Moreover, LSTM has three gate structures (inputting gate, forgetting gate, and outputting gate) and a Cell State’s hiding state. In LSTM, the inputting gate is responsible for processing the input of the current sequence position, whereas the forgetting gate controls whether the hidden cell state of the upper layer must be forgotten based on probability. The results of the forgetting gate and inputting gate will act on the cell state. Then, information from the previous sequence, the current sequence, and the cell state will be combined with the activation function and weights to obtain the output. Therefore, the model can better capture semantic information of peptide sequences and the prediction can more accurately select Bi-LSTM, as shown in **Supplementary Figure 1**.

To the best of our knowledge, our model is the first to determine the probability of each residue belonging to specific secondary structures by adding a linear layer with the softmax function behind the Bi-LSTM, after which the label with the highest probability can be obtained. However, this will ignore the correlation among secondary structures and decrease the prediction performance. Alternatively, we chose the CRF approach, which is widely used in named entity recognition to predict secondary structures, while exploring the context-related interactions between secondary structures and residue level contributions to all secondary structures.

CRFs consist of emission matrices including the probability of residues occupying different secondary structure states and transition matrices including the likelihood of transferring one secondary sub-structure state to another. During the training process, the model uses the forward and backward algorithms to infer the conditional probability of the secondary structures at each position of the sequence and finally predict the secondary structure by the scoring matrices. The specific calculation process is described below.

There are two kinds of feature functions. The first is referred to as the emission function, which is only related to the current position *i* in the peptide sequence:

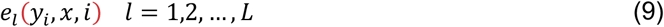

where *x* represents all residues of the peptide, *y_i_* represents the secondary structure at position *_i_*, and *L* indicates the number of all secondary structures.

The second function is defined in the context of secondary structures and is referred to as the transition function, which is related to the current structure *y_i_* and the previous structure y_i-1_:

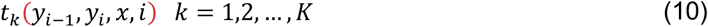

where *K* indicates the number of all permutations of two secondary structure states, which is 9 for 3-state secondary structures and 64 for 8-state secondary structures.

Assuming that we have *K_1_* transition functions and *K-_2_* emission functions, there are a total of *K_1_* + *K-_2_* feature functions. We then used the formula *f_k_*(*y_i-1_,y_i_,x,i*) to express them:

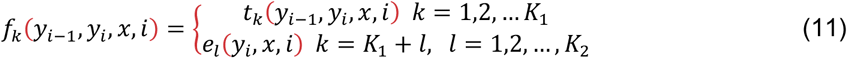

We also unified the weight coefficient *f_k_*(*y_i-1_,y_i_,x,i*) with w_k_:

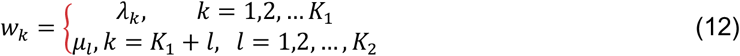

where *λ_k_* represents the weight coefficient of the k-th transition function and *μ_l_* represents the weight l-th coefficient of the emission function.

The parametric form is simplified as:

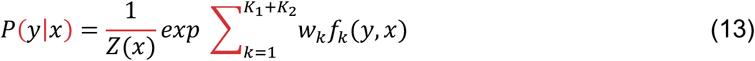

*Z*(*x*) is the normalization factor:

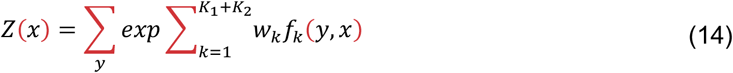

In the traditional CRF, we find that the only global transition matrix is easily affected by the noise from datasets, resulting in unstable prediction results. To solve this problem, we first arranged the outputs from Bi-LSTM into linear layers, transferring the outputs to local transition matrices with the same dimension as the global transition matrix. Then, we connected them to the global transition matrix, as using the fused transition matrices can improve the ability of our model to assess different datasets. The details of our CRF architecture are shown in **Supplementary Figure 2**.

### Model training and predicting process

#### Training process

We introduced the Bi-LSTM-CRF layer to fuse features and predict the secondary structure of peptides. In Bi-LSTM-CRF, the secondary structure label paths are constructed with the emission and transition matrices. The loss function of our model consists of two parts, the score of the real label path and the total score of all paths, with different secondary structure label combinations. The score of the real path should be the highest in all paths and the goal of our optimization is to minimize the gap between the predicted score and the real score.

If a certain path is a real path and the secondary structure label sequence is the correct prediction result, then it should have the highest score of all possible paths. According to the following loss function, the parameters of our model will be updated continuously with every iteration of the training process, making the ratio of the score of the real path to the total score larger.

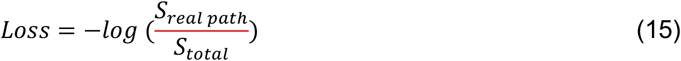

Assuming that the score of each possible path is *S_i_*, and there are *n* paths in total, then the total score of all paths is (where *e* is Euler number):

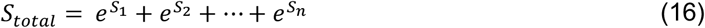

Next, the composition of *S_k_* can be expressed as follows:

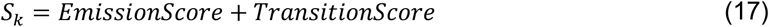

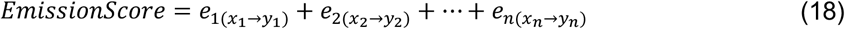

The *e*_*i*(*X_i_→y_i_*)_ is the score function resulting in a probability to predict the current residue *x_i_* as the secondary structure *y_i_*.

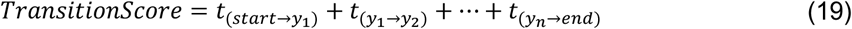

where *t*_*i*(*y_i_→y_j_*.)_ is the score function in support of generating the probability of transferring the secondary structure *y_i_* to *y_j_*.

#### Prediction process

In the prediction process, the Viterbi algorithm [37] is used to obtain the secondary structure prediction. The Viterbi algorithm is a dynamic programming algorithm that uses the start and end states and the recurrence formula to gain the secondary structure labels. The input of the Viterbi algorithm consists of *K* feature functions of the model, *K* weights related to the functions, the observation peptide sequence *x = (*x*_1_,*x*_2_,…,X_n_*), and the number of secondary structure states *m*. The output of this calculation is the optimal prediction secondary structure label sequence 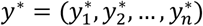. The details of the prediction process of the Viterbi algorithm are described below.

First, the start recursive algorithm is initialized as:

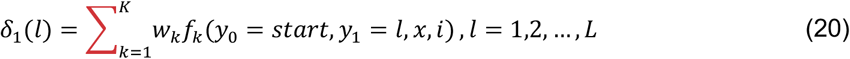

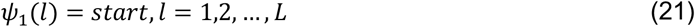

where *L* is the number of secondary structure labels.

For *i* = 1,2,…,*n* - 1, the recursion formula is performed as follows:

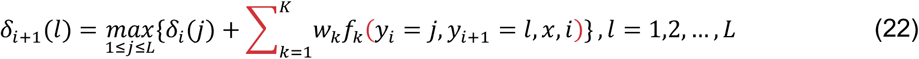

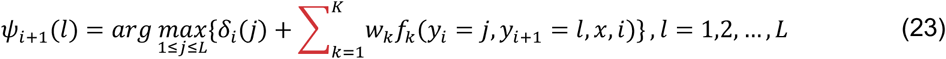

When the following condition occurs, program recursion is stopped:

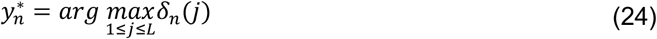

Through the backtracking algorithm, we obtain the final prediction structure:

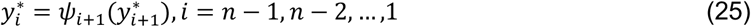

In the end, the prediction is:

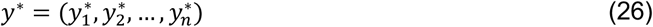

#### Performance metrics

The performance of our proposed PHAT is evaluated by the accuracy and SOV (segment overlap measure) for each secondary structure state. Acc_i_, F1-score_i_ {*i* represents the secondary structure element [H(Helix), E(Sheet) or C(Coil) for 3-state and H(alpha-helix), G(3_10_helix), I(π-helix), E(extended beta-strand), B(isolated beta-strand), T (turns), S (bend) and others (C) for 8-state]}, the accuracy in all states (hereinafter referred to as Acc), and SOV are calculated as follows:

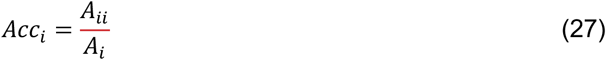

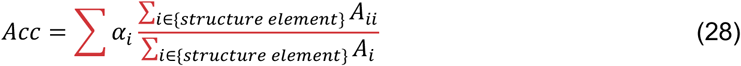

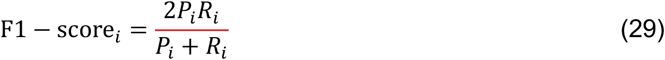

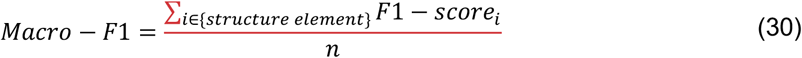

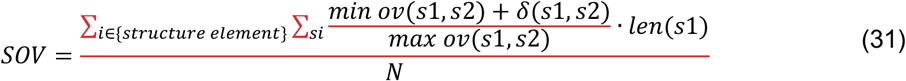

where *A_i_* is the sum of correctly predicted residues in each state; *A_ii_* is the number of correctly predicted residues in state i; *α_i_* is the proportion of state *i* in the entire test set; *P_i_*indicates the proportion of residues correctly predicted to be *i* among those predicted to be i; *R_i_* is the proportion of residues correctly predicted to be *i* among residues with the actual i; *s*1 and *s*2 are segments corresponding to actual and predicted secondary structures; *len*(*s*1) represents the number of residues defining the segment *s*1; max ov(*s*1,*s2*) is the maximum length overlap of *s*1 and *s*2 for which either of the segments has a residue in state *i*; min ov(*s*1,*s*2) represents the length overlapping s1 segments and *s*2 segments. *δ*(*s*1,*s*2) is calculated as follows:

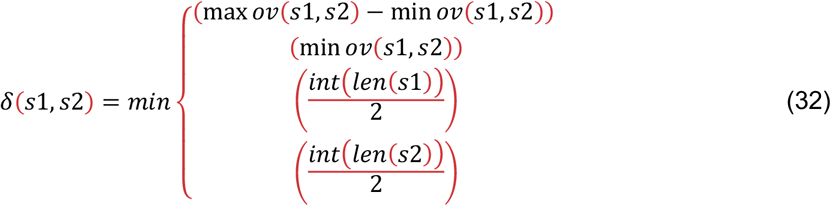

The normalization value *N* is a sum of *N*(*i*) over the entire set of conformational states:

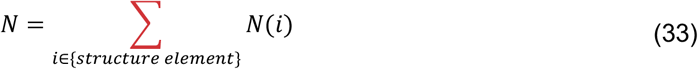

SOV was introduced because the segment overlap measure treats H, E, and C on an equal basis (eight-state assignment is the same). There are no arbitrary cutoffs on segment length, thus ensuring a consecutive and threshold-free assessment of prediction accuracy.

## Results

### PHAT outperforms existing methods when analyzing an independent testing set

To evaluate the performance of the proposed PHAT model, we compared it with four state-of-the-art methods: PROTEUS2 [14], RaptorX [16], Jpred [12], and PSSP-MVIRT [19]. The first three were designed for protein secondary structure prediction whereas the other is for peptide secondary structure prediction. To ensure a fair comparison, the models were executed and evaluated using the same independent test set. As shown in **Supplementary Table 3**, PHAT achieved the best performance among all of the tested methods, with an Acc of 84.07%, Acc_H_ of 89.08%, Acc_E_ of 71.76%, Acc_c_ of 80.66%, and SOV of 79.78%. Specifically, compared to other existing methods, our method delivered 1.39% to 19.26% higher SOV values (see **Figure 2A and Supplementary Table 3**), which is an important metric at the segment level and evaluates the overall performance of the methods. The superior SOV performance of our proposed model might be related to the context information of the peptide sequences extracted by the Bi-LSTM-CRF layer and multi-scale features captured by the hypergraph multi-head attention network. Furthermore, all methods exhibited a relatively low accuracy in the prediction of the structural state E compared to the other two states (H and C). This was due to the low proportion of E in the dataset (see **Figure 2B and Supplementary Table 1**). Therefore, the existing models capture more information for the H and C states, rather than E, during model training. Nevertheless, our PHAT achieved the highest accuracy at E among all of the evaluated methods. This was likely because our multi-head attention mechanism is capable of capturing a more informative structural representation of E. Additionally, the comparison results in the dataset of the eight-state secondary structure shown in **Supplementary Table 9** also demonstrate the outstanding performance of our method. Therefore, we concluded that our method is more effective than Jpred, PSSP-MVIRT, PROTEUS2, and RaptorX in the prediction of peptide secondary structures, especially for Acc_E_, Acc, and SOV.

**Figure 2.**
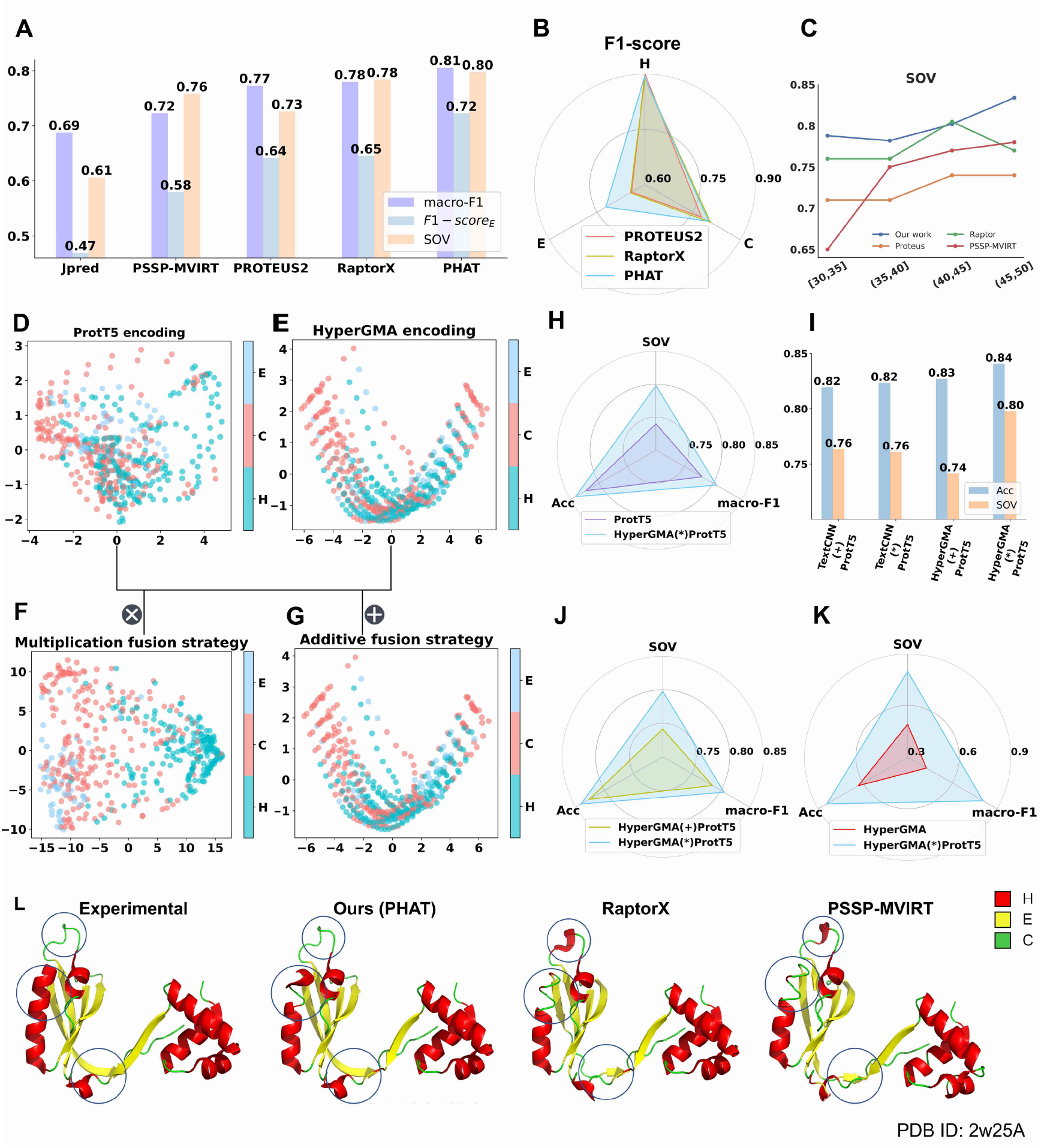
The performances of our method and existing methods on independent test subsets, comparison of different encoding strategies, and visualization of different methods on one peptide: (**A**) SOV, macro-F1, and F1-scoreH are used as the evaluation metrics; (**B**) F1-scores under three sub-structures are used as the evaluation metrics. (**C**)SOV of four methods at the different length intervals. (**D–G**) represent PCA visualization results of individual features of ProtT5, HyperGMA, and the fusion features in multiplication or additive respectively; (**H, J, K**)represent the comparison between multiplication fusion strategy and other three strategies. (**I**)represents performance comparison between HyperGMA and TextCNN. (**L**)The visualization of predictions by our method and other two methods for the peptide with PDB ID: 2w25.

### Length preference investigation for peptide secondary structure prediction

Previous studies have demonstrated that the functionality of peptides (e.g., affinity) is easily affected by the length of sequences, with most bioactive peptides being normally less than 40 residues long[19, 38, 39]. To investigate if our model had length biases for peptide secondary structure prediction, we further explored whether peptide length affected the performance of our model. We first divided the test set into four subsets with different length intervals ([30, 35], (35, 40], (40, 45], and (45, 50]), then separately evaluated our model and existing methods using the subsets. **Figure 2C** and **Supplementary Figure 3** show the SOV, Acc, and F1-score of the different methods for the prediction of peptide secondary structures using the aforementioned subsets. As illustrated in **Supplementary Figure 3**, the performance of all of the tested methods clearly decreased as the length of the sequences declined, which indicates that shorter sequences are more difficult to predict as their contextual information is less. Furthermore, as illustrated in **Figure 2C**, the SOV score of our method was higher than that of the other methods in almost all ranges of peptide sequence lengths. Particularly, our PHAT model exhibited an outstanding performance, with average Acc, SOV, and F1-score values up to 7.02%, 6.21%, and 3.33% higher than the runner-up PSSP-MVIRT in different sequence length intervals. These results demonstrate that our method is better at the prediction of shorter peptides.

### Exploration of the optimal architecture of our model

To investigate the performances of our model using different encoding strategies, we compared the prediction results of different encoding strategies including the two individual feature extractors (HyperGMA and ProtT5) and their different fusion combinations. **Supplementary Table 4** shows that our final element-wise multiplication strategy achieves an Acc of 84.07%, Acc_H_ of 89.08%, Acc_E_ of 71.76%, Acc_c_ of 80.66%, and SOV of 79.78%, outperforming the Acc and SOV of ProtT5 by 1.77% and 5.79% and the fused extractor in the additive strategy by 1.36% and 5.64%, respectively. Furthermore, although ProtT5 performed better than HyperGMA, the model performed better than the individual extractors and the fused extractor in the additive strategy after fusing the features from HyperGMA and ProtT5 with the element-wise multiplication fusion strategy. This indicated that the different information is complementary to each other in the fusion strategy, thus effectively improving the predictive performance of the model. Moreover, it can be seen from **Figure 2H-2K** that the element-wise multiplication fusion strategy of HyperGMA and ProtT5 achieved better performance than the fusion strategies of TextCNN and ProtT5 in terms of Acc and SOV.

To further illustrate the effect of different encoding strategies more intuitively, we visualized the distribution of feature representations in the test set, which reveals the discriminability of features for distinguishing different secondary sub-structure states through dimension reduction. In the principal component analysis (PCA) [40] in **Figure 2D–2G**, each point represents a site in the peptide sequence and different colors are used to distinguish the Helix (H), Strand (E), and Coil (C) secondary structures. Compared with the two fusion strategies, the distribution of the site samples belonging to different classes in the feature space from the individual ProtT5 and HyperGMA are almost connected, making it difficult to distinguish the region for each secondary sub-structure class. Regarding the two fusion strategies, the site samples of three classes are more clearly distributed in the feature space of the multiplication fusion strategy (**Figure 2F**) than in the feature space of the additive fusion strategy (**Figure 2G**). Furthermore, to avoid biases between different dimension reduction methods, we also applied another non-linear method T-SNE [41] for dimension reduction, and similar results can be seen in **Supplementary Figure 3**. In conclusion, our results demonstrate that our PHAT model with the multiplication fusion strategy can capture more discriminative and high-quality features.

### The PHAT model has good interpretability in terms of extracting multi-scale features and making classifications

To verify the effect of the Bi-LSTM-CRF layer in our model, we compared the performance of our model under two training strategies (Cross Entropy loss function and Bi-LSTM-CRF), and the results are shown in **Supplementary Table 5**. Clearly, our model with Bi-LSTM-CRF layer performed better (especially in terms of SOV) than the model using the Cross-Entropy loss function. To explain how the Bi-LSTM-CRF efficiently predicts the secondary structure at each site in the peptide sequence, we randomly selected and predicted the secondary structures of the peptide sequence with PDB ID 1edm chain B (Protein Data Bank Identity). Afterward, we chose several sites of this peptide and visualized the corresponding weights of the transition matrix and emission matrices from our model in **Figure 3A**. As illustrated in **Figure 3A**, the secondary structure labels corresponding to the highest values in the emission matrices match the real secondary structures of the residues. Moreover, the probability of transferring the labels of the current residues to the real labels of the adjacent residues was the highest in the transition matrices.

**Figure 3.**
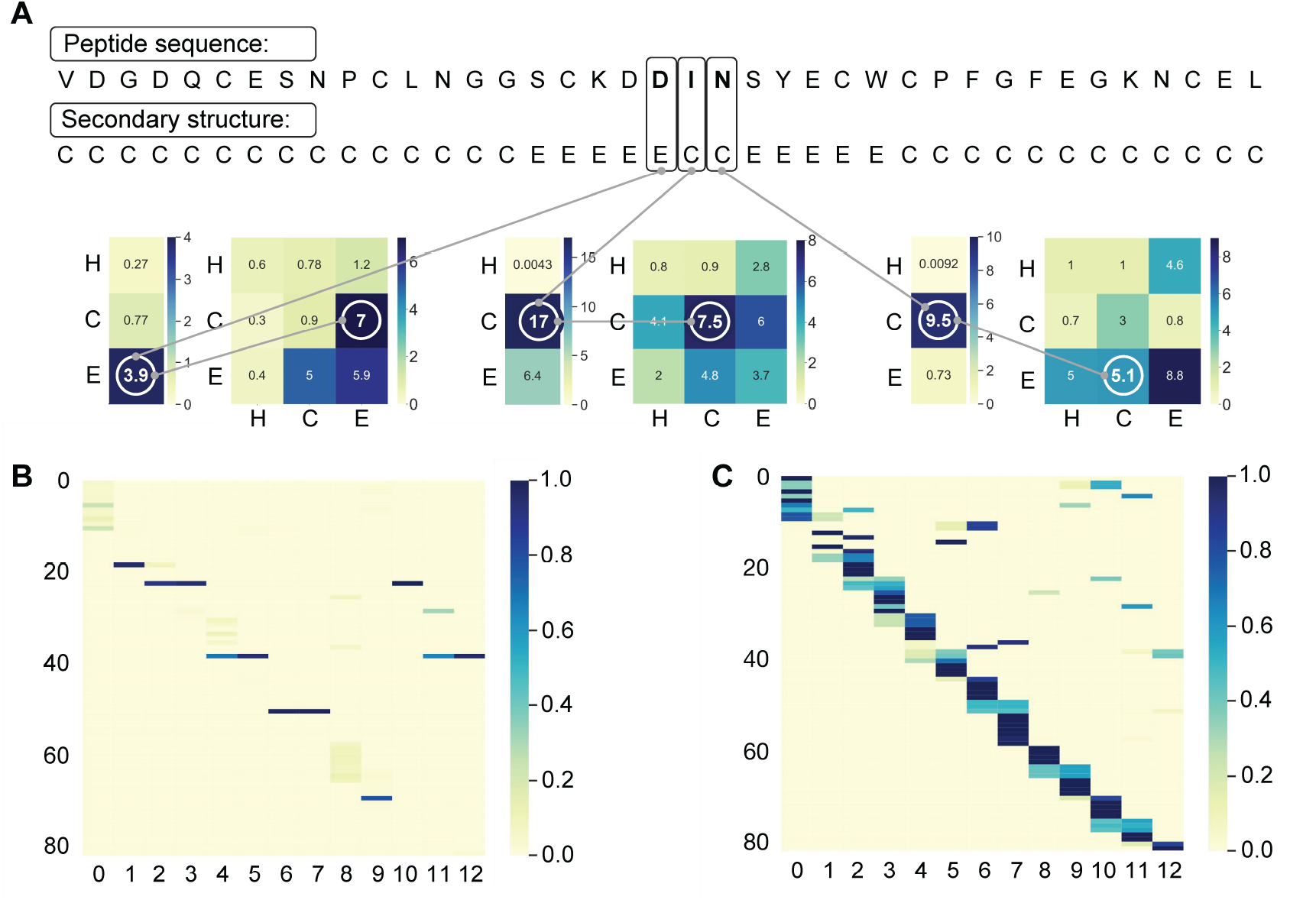
The Interpretability of our model. (**A**) Visualization of the weights of transition matrix and emission matrix in Bi-LSTM-CRF layer. The emission matrix and transition matrix are calculated by our model. The emission matrix shows the possibilities of current site in different classes and the transition matrix indicates the possibility of the secondary structure transformation in adjacent positions. (**B-C**) Visualization of the attention matrices in hypergraph multi-head attention network, where **B** represents the attention of peptide fragments to residue groups and **C** represents the attention of residue groups to peptide fragments. Darker color means stronger attention.

To further explore the role HyperGMA of in our model, we visualized and analyzed the attention matrices from HyperGMA in **Figure 3**. The HyperGMA includes two main steps, the residue group level attention encoding and the peptide fragment level attention encoding. In the first step, the feature representations of peptide fragments are aggregated from the contained residue groups through the residue group level multi-head attention mechanism. The contribution of each residue group to corresponding peptide fragments is shown in **Figure 3B**. Moreover, **Figure 3C** illustrates that the peptide fragments are more likely to reflect the characteristics of specific residue groups, meaning that the peptide fragments are more strongly influenced by local information. In the second step, the feature representation of the residue group is encoded by the peptide fragments where it exists through the fragment level attention mechanism. The contribution of the peptide fragment to corresponding residue groups is shown in **Figure 3C**, which indicates that a given residue group can aggregate the information from different fragments where it exists. Therefore, our model can better capture the local and global information by collecting secondary structure information at the residue group level and peptide fragment level using HyperGMA.

### Application of our PHAT model in three peptide related downstream tasks

Several experiments were conducted to verify that the secondary structures predicted by our method can be useful for downstream tasks. **Figure 4A–4C** shows the results of prediction of peptide toxicity, prediction of T-cell receptor interactions with MHC-peptide complexes, and prediction of protein-peptide binding sites, respectively. In **Figure 4**, it can be seen that when fused with the structure predictions of our PHAT model, the evaluated methods (ATSE, NetTCR-2.0, and PepBCL) achieve higher performance in terms of most metrics than without the PHAT predictions. Similar results were observed with the methods fused with structure predictions from PROTEUS2 and PSSP-MVIRT in the corresponding task.

**Figure 4.**
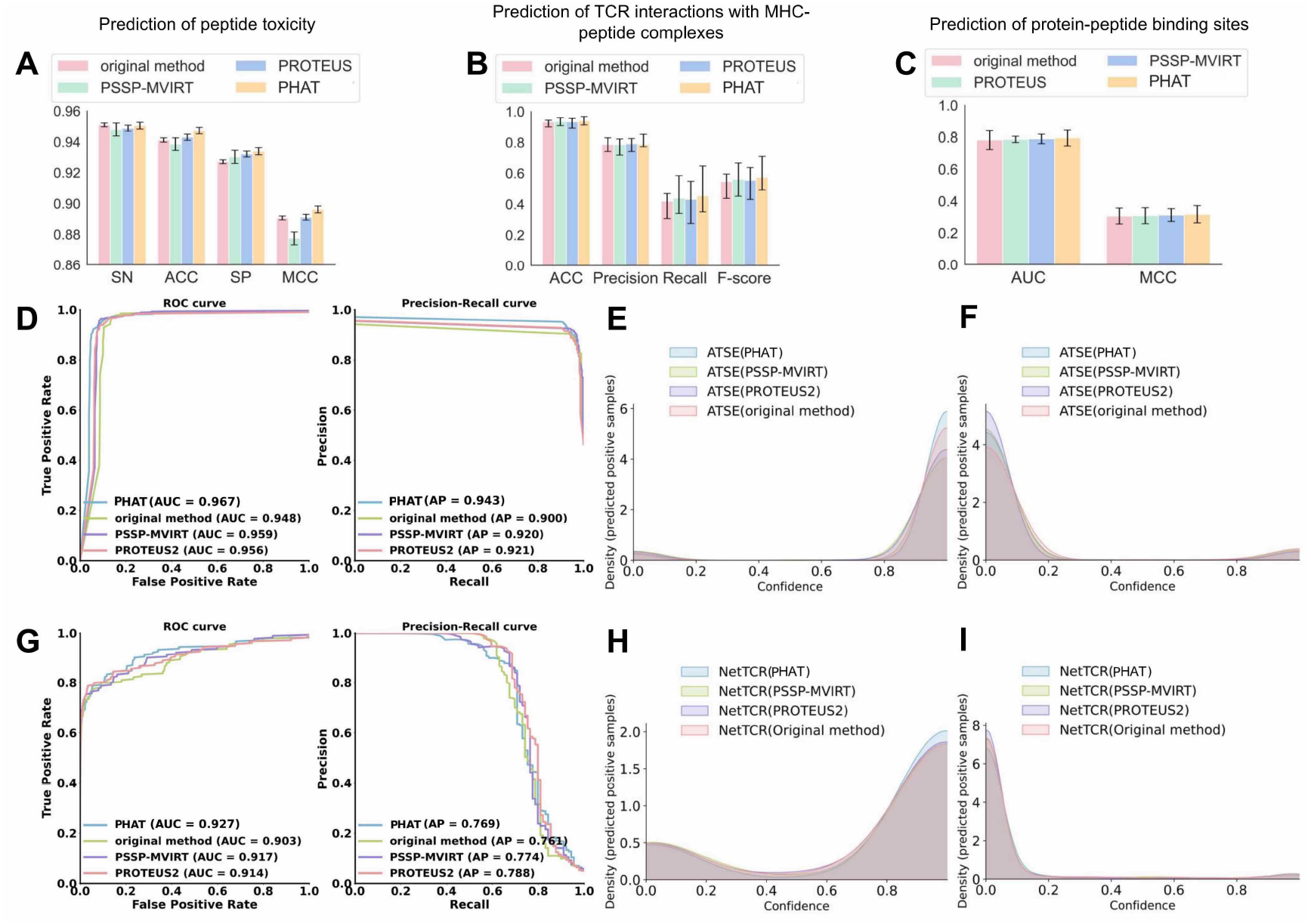
Comparative results for three downstream tasks. (**A**) shows the results on the task of prediction of peptide toxicity. (**B**) shows the results on the task of prediction of T-cell receptor interactions with MHC-peptide complexes. (**C**) shows the results on the task of prediction of protein-peptide binding sites. (**D**) shows ROC curve and Precision-Recall cure of comparison experiment in ATSE. (**E**) and (**F**) show density of positive and negative examples under different confidence in prediction of peptide toxicity. (**G**) shows ROC curve and Precision-Recall cure of comparison experiment in NetTCR-2.0. (**H**) and (**I**) show density of positive and negative examples under different confidence in prediction of TCR interactions with MHC-peptide complexes.

### PHAT has an outstanding performance for aiding in predicting peptide toxicity

We first used the methods (PSSP-MVIRT, PROTEUS2, and PHAT) to predict the secondary structures of the dataset in ATSE [26], a peptide toxicity predictor, and add the secondary structures from the three methods to ATSE. As shown in **Figure 4D** and **Supplementary Table 6**, ATSE with our PHAT model achieved an SN of 95.06%, SP of 93.4%, Acc of 94.74%, MCC of 89.62%, AUC of 96.7% (the definition of these metrics can be found in Supplementary metrics), which constituted a 0.17%, 0.18%, 0.43%, 0.5%, and 1.1% higher performance than ATSE with PROTEUS2, and a 0.25%, 0.37%, 0.88%, 1.87%, and 0.8% higher performance than ATSE with PSSP-MVIRT, respectively. Additionally, **Figure 4E–4F** shows PHAT had an outstanding performance for the prediction and classification of ATSE, and there was also a general improvement over the original method. These results demonstrate the efficiency of our model to predict secondary structures to assist in peptide toxicity prediction. Particularly, the higher SOV of our method reveals that our model can more accurately capture the integrity and continuity of secondary structures, which may explain the superior performance of our method.

Secondary structure is an important determinant of toxicity [42]. However, few studies have used the secondary structure of peptides to predict peptide toxicity. Predicting the secondary structures of peptides by various methods can compensate for these limitations and build a bridge between peptide secondary structure and peptide toxicity.

### PHAT achieves superior performance for the prediction of T-cell receptor interactions with MHC-peptide complexes

Our prediction of the secondary structure of peptides can also be applied to the study of T-cell receptor interactions with MHC-peptide complexes. Here, we used the NetTCR-2.0 method [27], which has a CNN architecture, to predict the interactions between the α/β TCR and MHC-peptide sequences and assess the effect of adding secondary structures predicted from the three methods (PSSP-MVIRT, PROTEUS2, and our PHAT). As indicated in **Figure 4G** and **Supplementary Table 7**, analysis of the NetTCR-2.0 dataset with PHAT achieved an average Acc of 94.04%, a precision of 45.54%, a recall of 78.6%, an F1-score of 57.29%, and an AUC of 92.7%, which was higher than the original method by 0.61%, 3.52%, 2.61%, and 2.4%, respectively. Furthermore, our model outperformed the Acc, Precision, F1-score, and AUC of PSSP-MVIRT by 0.38%, 1.61%, 1.28%, and 1%, as well as PROTEUS2 by 0.59%, 2.29%, 1.82%, and 1.3%, respectively. Moreover, **Figure 4H–4I**shows that PHAT achieved a better prediction of NetTCR-2.0 classification.

Additionally, we found that two groups of α/β TCR sequences, which have similar sequences but different secondary structures, cannot be classified correctly using NetTCR-2.0 without adding secondary structures. Fortunately, they were accurately predicted after introducing the secondary structure features from our PHAT model. In **Supplementary Figure 4**, we visualized the secondary structures of the two peptide sequences predicted by our method. Therefore, our findings demonstrated that the secondary structures predicted by our method provide useful biochemical information and improve the performance of NetTCR-2.0. In conclusion, the above results can prove that our prediction of peptide secondary structures has a positive effect on promoting the accuracy of TCR tasks and provide a new direction for TCR research.

### PHAT exhibited competitive performance for assisting in the prediction of protein-peptide binding sites

Protein-peptide interactions are involved in various fundamental cellular functions and are crucial for designing new peptide drugs. To explore the effect of the secondary structures from our model in the prediction of protein-peptide binding sites, comparison experiments with the PepBCL model were conducted [43]. Specifically, we first combined our structure predictions with the features from the PepBCL model. Then, protein-peptide binding site predictions were conducted based on a random forest machine learning method [44]. In a previous study that used the PepBCL model [36], the secondary structure from SPOT-1D-Single was introduced to generate structural features, which we generated in the same way. In this context, the efficiency of our prediction can be verified by comparing secondary structures from several different sources (**Supplementary Table 8**). Our findings indicated that the application of peptide secondary structures predicted by our PHAT achieves significantly better performance than other methods. Some researchers have already incorporated secondary structures into their predictions. Moreover, the prediction of more accurate and continuous secondary structures may enhance the efficiency of site mining. As illustrated in **Figure 4C**, the features from PepBCL combined with the prediction of PHAT can achieve higher AUC and MCC than using peptide secondary structures from other methods.

### The visualization of two cases demonstrated that our proposed PHAT method performs better than existing methods

To intuitively assess the performance of existing methods, we first randomly selected two peptide chains (PDB ID: 2w25A and 1ejbA) with experimental secondary structures, and applied different methods (PHAT, RaptorX, PSSP-MVIRT, PROTEUS2, and Jpred) for the prediction of the secondary structure of two peptides. As illustrated in **Figure 2L and Supplementary Figure 5**, the secondary structures from different methods were mapped into the tertiary structures, where the red area represents Helix (H), the yellow area represents Strand (E), and the green area represents Coil (C). The differences between the structures predicted by our method and the experimental ones were smaller than those of the predictions of the other methods described above. In **Figure 2L**, our model achieved more correct Helix (H) and Strand (E) predictions, whereas the other methods were more likely to identify the Helix (H) and Strand (E) structures as a Coil (C). Furthermore, in **Supplementary Figure 5**, the other four methods (RaptorX, PSSP-MVIRT, PROTEUS2, and Jpred) tended to predict the Coil (C) as Helix (H), whereas our method made more correct predictions in local consecutive sequence regions. In conclusion, our method can achieve better performance in terms of continuity and accuracy compared to the existing methods.

### The proposed PHAT model facilitates the construction of 3-D peptide structures

To explore the potential of PHAT in capturing 3-D structure information of peptides, we used our model to predict the distance map and contact map matrices, which is an essential process in protein 3-D structure prediction. The workflow of exploration is shown in **Figure 5A**. Specifically, our PHAT model was first trained using a secondary structure dataset to capture the 2-D structure information of the peptide. Then, a fully connected network was added to our model and fine-tuned using the contact map dataset (the details are shown in **Supplementary Table 10**) to obtain the distance information of the 3-D structure. Next, we calculated the distance of each amino acid pair to construct the distance map and contact map of the peptide sequence. Compared with the experimental results from the test set, our model achieved an average variation of less than 1 *Å* for each amino acid pair in terms of distance map prediction. To intuitively assess the performance of our model, we visualized and compared our predictions with the state-of-the-art method trRosetta [45–47] based on the experimental results from a randomly selected peptide with PDB ID 7ve4 (**Figure 5B**). Our predicted contact map is more accurate in terms of contacting amino acid pairs than the one obtained with trRosetta. Additionally, our predicted distance map is closer to the experimental result than the trRosetta-generated map, indicating that our model can more accurately capture the distance between amino acids. With our predicted contact maps and distance maps, the 3-D structures of corresponding peptides can be reconstructed more realistically by folding algorithms [48–50]. In this case, we extended our prediction of the secondary structure to the contact map and distance map, thus aiding in the prediction of the peptide 3-D structure. Therefore, our PHAT model has the potential to promote the development of therapeutic molecules against various diseases, as well as the design of functional peptides [40].

**Figure 5.**
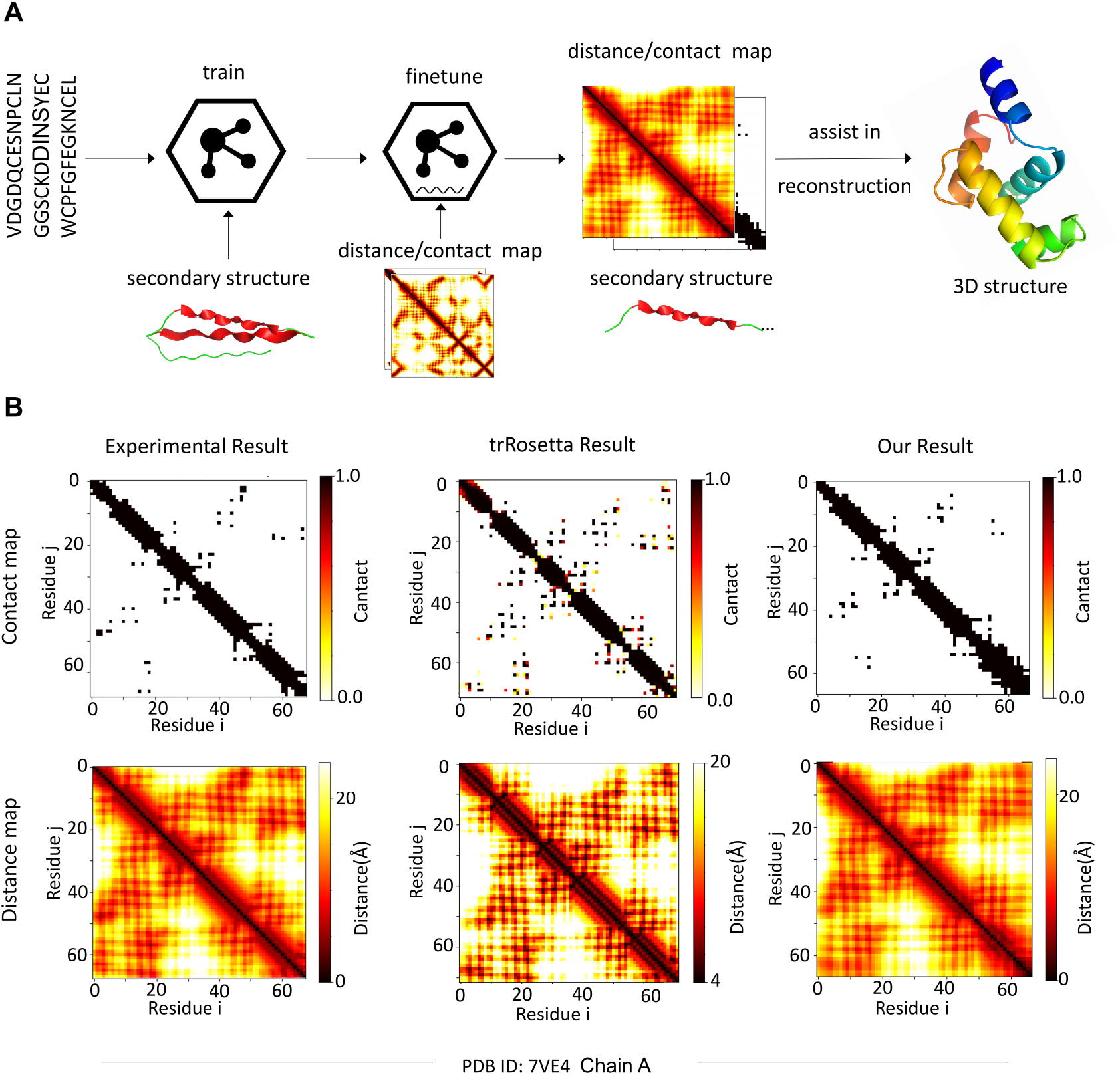
The exploration in constructing 3-D structure of peptide with our method. (**A**) The workflow of assisting in building 3-D peptide structure with our predicted contact and distance map matrices. (**B**) The visualization of contact map and distance map matrices of experimental results, trRosetta prediction and our prediction for the peptide with PDB ID: 7ve4.

## Discussion and Conclusion

In this study, we developed PHAT, a deep learning-based method for peptide secondary structure prediction, and systematically evaluated it using benchmark datasets. Compared with other methods designed for protein secondary structure prediction, our model achieved superior performance in most metrics, especially Acc_E_ and SOV. The conventional methods designed for the prediction of protein structure might be biased toward extracting long-distance dependence within protein sequences with hundreds of residues. However, the peptides in our dataset are significantly shorter than most proteins, and therefore the neighborhood information in peptides may not be easily captured by these methods. In contrast, our method can capture more contextual information of peptide sequences through the hypergraph multi-head attention network, and can thus make more correct predictions in local consecutive sequence regions, as demonstrated by the visualization of our predictions for two peptides (PDB ID: 2w25A and 1ejbA).

Similar results can be seen when comparing the peptide-specific secondary structure predictors (e.g., PSSP-MVIRT) with our method. This is likely because previous methods designed for peptides focus more on neighborhood information of peptide residues and therefore tend to ignore long-term information. In contrast, in addition to being capable of capturing contextual information, our method can obtain long-term and bio-semantic knowledge for peptide sequences by using ProtT5, a model pre-trained with millions of protein sequences, thus achieving a good prediction performance. The peptide length preference experiments for secondary structure prediction illustrated that although the prediction performance of the tested methods decreased as the length of the sequences declined, our method achieved better performance than other existing methods when analyzing shorter peptide sequences. This indicated that our model can integrate contextual information and long-term knowledge to make predictions.

Moreover, to reveal the feature extraction and prediction mechanisms of our PHAT model, we visualized matrices of a hypergraph multi-head attention network (HyperGMA) and Bi-LSTM-CRF, which provide good interpretability while achieving an outstanding prediction performance. Specifically, the visualization of attention matrices in HyperGMA demonstrated that our model can effectively capture the local and global features of peptides at the residue group-level and the peptide fragment-level, thus providing insights into its attention mechanisms. Similarly, the visualization of the classification layer in Bi-LSTM-CRF illustrates that CRFs can guide our model to efficiently predict the secondary structure for each site in the peptide sequences.

Furthermore, to verify the accuracy of the secondary structures predicted by our model in downstream tasks, we applied our predicted structural information to the prediction of peptide toxicity, T-cell receptor interactions with MHC-peptide complexes, and identification of protein-peptide binding sites. Using the secondary structures predicted by our model enhanced the performances of these tasks, which indicated that our predicted structural information can assist in predicting more accurate properties and is complementary to sequential and evolutionary features in peptide-related downstream tasks. Additionally, to explore the potential of PHAT in capturing 3-D structural information of peptides, we applied our model to predict distance map and contact map matrices and achieved an outstanding performance, thus demonstrating that our model can help in the reconstruction of peptide 3-D structures. We also developed an online service (the workflow is shown in **Figure 1D**) to implement our PHAT, thus saving researchers the need to write programs or scripts. We hope that this online tool will be helpful to the research community.

Although our PHAT model achieves improved performances for predicting peptide secondary structure, there is still room for improvement. For example, PHAT is meant to be used for general peptide secondary structure prediction, and therefore we focused particularly on sequences with lengths <50. However, for datasets with peptide sequences longer than 50, we cannot ensure that our method will have the same performance. Moreover, when interacting with other targets (e.g., protein, DNA, RNA, *etc.*), peptide sequences remain the same, but the secondary structure of the peptides may change considerably. However, our PHAT makes its predictions based on the sequence patterns and thus cannot make adjustments to account for potential molecular interactions. Therefore, we are planning to incorporate additional data such as interaction information with other targets to further improve the prediction of peptide secondary structures in different interacting scenarios.

## Data Availability

The authors declare that the data supporting the findings of this study are available within the article and its supplementary information files. Besides, the benchmarking datasets and our source code were also available for downloading at http://inner.wei-group.net/PSSPHAT/.

## Funding

The work was supported by the Natural Science Foundation of China (Nos. 62071278 and 62072329), and Natural Science Foundation of Shandong Province (ZR2020ZD35).

## Conflict of Interest

The authors declare that they have no competing interests.

## Notes

### Competing Interest Statement

The authors have declared no competing interest.

### Summary of Updates

The title and abstract updated to clarify the highlights of our model

