## Supplementary Materials for "Explainable deep graph learning accurately modeling the peptide secondary structure prediction"

### Supplementary Tables

**Supplementary Table 1. Summary of the datasets for peptide secondary structures of three states.**

| Datasets | Structural states | | | Sequence number |
|---|---|---|---|---|
| | H | E | C | |
| Training set | 33,455 | 16,901 | 29,177 | 1,028 |
| Testing set | 5,294 | 1,119 | 3,733 | 257 |

**Supplementary Table 2. Summary of the datasets for peptide secondary structures of eight states.**

| Datasets | Structural states | | | | | | | | Sequence number |
|---|---|---|---|---|---|---|---|---|---|
| | H | G | I | E | B | T | S | C | |
| Training set | 48,132 | 7,262 | 35 | 49,205 | 2,994 | 24,006 | 22,394 | 46,940 | 1,018 |
| Testing set | 1,616 | 150 | 0 | 1,089 | 86 | 707 | 513 | 966 | 42 |

Note: We randomly selected 1,018 sequences out of all 1,060 sequences as the training set and the remaining 42 sequences as our test set.

**Supplementary Table 3. The experimental results of all the methods on independent test set**

| Method | $\text{Observed}_j$ | Predicted | | | $\text{Acc}_j$ (%) | Acc (%) | SOV (%) |
|---|---|---|---|---|---|---|---|
| | | H | E | C | | | |
| Jpred | H | 4,195 | 146 | 953 | 79.24 | 78.05 | 60.62 |
| | E | 64 | 467 | 588 | 52.54 | | |
| | C | 337 | 259 | 3,136 | 84.03 | | |
| PSSP-MVIRT | H | 4,773 | 99 | 422 | 90.16 | 78.50 | 75.81 |
| | E | 139 | 636 | 344 | 56.84 | | |
| | C | 836 | 341 | 2,556 | 68.47 | | |
| PROTEUS2 | H | 4,656 | 81 | 557 | 87.95 | 82.45 | 72.61 |
| | E | 34 | 770 | 315 | 68.81 | | |
| | C | 364 | 430 | 2,913 | 78.73 | | |
| RaptorX | H | 4,493 | 142 | 659 | 84.87 | 82.98 | 78.39 |
| | E | 27 | 693 | 399 | 61.93 | | |
| | C | 306 | 194 | 3,233 | 86.61 | | |

| | | | | | | | |
|---|---|---|---|---|---|---|---|
| | H | 4,716 | 61 | 517 | 89.08 | | |
| PHAT | E | 49 | 803 | 267 | **71.76** | **84.07** | **79.78** |
| | C | 482 | 240 | 3,011 | 80.66 | | |

**Supplementary Table 4. Results of the models with different encoding strategies.**

| Method | Observed$_j$ | Predicted | | | $Acc_j$ | Acc | SOV |
|---|---|---|---|---|---|---|---|
| | | H | E | C | (%) | (%) | (%) |
| | H | 4,698 | 294 | 302 | 88.74 | | |
| HyperGMA | E | 191 | 192 | 736 | 65.77 | 63.04 | 49.08 |
| | C | 1,256 | 1,515 | 962 | 25.77 | | |
| | H | 4,814 | 133 | 347 | 90.93 | | |
| ProtT5 | E | 68 | 873 | 178 | 78.02 | 82.30 | 73.99 |
| | C | 629 | 440 | 2,664 | 71.36 | | |
| | H | 4,726 | 73 | 495 | 89.27 | | |
| HyperGMA(+)ProtT5 | E | 66 | 776 | 277 | 69.35 | 82.71 | 74.14 |
| | C | 553 | 290 | 2,890 | 77.42 | | |
| | H | 4729 | 89 | 476 | 89.32 | | |
| TextCNN(+)ProtT5 | E | 64 | 800 | 255 | 71.49 | 81.95 | 76.34 |
| | C | 606 | 341 | 2786 | 74.63 | | |
| | H | 4548 | 58 | 688 | 85.91 | | |
| TextCNN(*)ProtT5 | E | 66 | 679 | 374 | 60.68 | 82.33 | 76.08 |
| | C | 442 | 165 | 3126 | 83.74 | | |
| | H | 4,716 | 61 | 517 | 89.08 | | |
| HyperGMA(*)ProtT5 | E | 49 | 803 | 267 | 71.76 | **84.07** | **79.78** |
| | C | 482 | 240 | 3,011 | 80.66 | | |

Note: (+) represents the fusion of encoding features with the element-wise multiplication strategy, and (*) represents the fusion of encoding features with the element-wise additive strategy.

**Supplementary Table 5. Results of our model with different training strategies.**

| Method | Observed$_j$ | Predicted | | | $Acc_j$ | Acc | SOV |
|---|---|---|---|---|---|---|---|
| | | H | E | C | (%) | (%) | (%) |
| | H | 4,723 | 91 | 480 | 89.21 | | |
| Cross Entropy loss function | E | 63 | 801 | 255 | 71.58 | 83.24 | 77.32 |
| | C | 537 | 274 | 2,922 | 78.27 | | |
| | H | 4,716 | 61 | 517 | 89.08 | | |
| CRF score function | E | 49 | 803 | 267 | 71.76 | 84.07 | 79.78 |
| | C | 482 | 240 | 3,011 | 80.66 | | |

**Supplementary Table 6. The results of comparison in prediction of peptide toxicity.**

| Method | SN | SP | FDR | FPR | Acc | MCC |
|---|---|---|---|---|---|---|

| | | | | | | |
|---|---|---|---|---|---|---|
| ATSE (original method) | 95.11% (+0.12%, -0.12%) | 92.72% (+0.12%, -0.13%) | 8.72% (+0.13%, -0.13%) | 7.81% (+0.13%, -0.13%) | 94.13% (+0.14%, -0.13%) | 89.07% (+0.12%, -0.14%) |
| ATSE (PSSP-MVIRT) | 94.81% (+0.42%, -0.40%) | 93.03% (+0.42%, -0.44%) | 8.62% (+0.40%, -0.43%) | 7.99% (+0.41%, -0.41%) | 93.86% (+0.41%, -0.41%) | 87.75% (+0.40%, -0.44%) |
| ATSE (PROTEUS2) | 94.89% (+0.20%, -0.21%) | 93.22% (+0.18%, -0.20%) | 7.90% (+0.19%. -0.19%) | 7.24% (+0.21%, -0.20%) | 94.31% (+0.20%, -0.20%) | 89.12% (+0.19%, -0.18%) |
| ATSE (PHAT) | 95.06% (+0.20%, -0.21%) | 93.4% (+0.22%, -0.23%) | 8.51% (+0.21%. -0.21%) | 7.53% (+0.20%, -0.22%) | 94.74% (+0.20%, -0.20%) | 89.62% (+0.20%, -0.24%) |

Note: We report the average after performing each experiment 20 times by splitting the data set for other methods based on the data set of ATSE.

**Supplementary Table 7. The results of comparison in prediction of T-cell receptor interactions with MHC-peptide complexes.**

| Method | Acc | Precision | Recall | F1-score |
|---|---|---|---|---|
| NetTCR-2.0 (original method) | 93.43% (+1.07%, -3.23%) | 42.03% (+4.98%, -11.23%) | 78.67% (+4.44%, -4.47%) | 54.68% (+4.93, -10.78%) |
| NetTCR-2.0 (PROTEUS2) | 93.45% (+2.25%, -4.05%) | 43.25% (+11.55%, -15.75%) | 79.21% (+3.39%, -5.01%) | 55.47% (+8.33%, -12.27%) |
| NetTCR-2.0 (PSSP-MVIRT) | 93.66% (+2.44%, -2.65%) | 43.93% (+14.57, -9.83%) | 78.83% (+3.7%, -6.7%) | 56.01% (+10.69%, -10.71%) |
| NetTCR-2.0 (PHAT) | 94.04% (+2.76%, -2.54%) | 45.54% (+19.26, -10.44%) | 78.6% (+6.57%,1. 53%) | 57.29% (+13.81%, -7.99%) |

Note: We report the average after performing each experiment 20 times by splitting the data set for other methods based on the data set of NetTCR-2.0.

**Supplementary Table 8. The results of comparison in prediction of protein-peptide binding sites.**

| Method | AUC | MCC |
|---|---|---|
| PepBCL (SPOT-1D-Single) | 78.6% | 35.7% |
| PepBCL (PROTEUS2) | 79.02% | 31.3% |
| PepBCL (PSSP-MVIRT) | 78.7% | 30.9% |
| PepBCL (PHAT) | 79.6% | 36.0% |

Note: We report the average after performing each experiment 20 times by splitting the data set for other methods based on the data set of PepBCL.

**Supplementary Table 9. The experimental results of PHAT and SSpro8 on independent test set for eight states.**

| Method | $Acc_j$(%) | | | | | | | | Acc (%) | SOV (%) |
|---|---|---|---|---|---|---|---|---|---|---|
| | H | G | I | E | B | T | S | C | | |
| PHAT | 78.06 | 67.78 | 0 | 77.55 | 63.21 | 76.10 | 68.32 | 74.37 | 75.49 | 76.11 |
| SSpro8 | 79.52 | 66.74 | 0 | 79.19 | 62.45 | 69.47 | 70.18 | 76.33 | 75.19 | 73.62 |

**Supplementary Table 10. Summary of the datasets for peptide distance/contact map**

| Datasets | Max length | Min length | Average length | Sequence number |
|---|---|---|---|---|
| Training set | 100 | 31 | 73 | 2715 |
| Testing set | 99 | 30 | 72 | 200 |

Note: The peptide sequences are from SCRATCH-1D and the corresponding structures are extracted from Protein Data Bank.

### Supplementary Figures

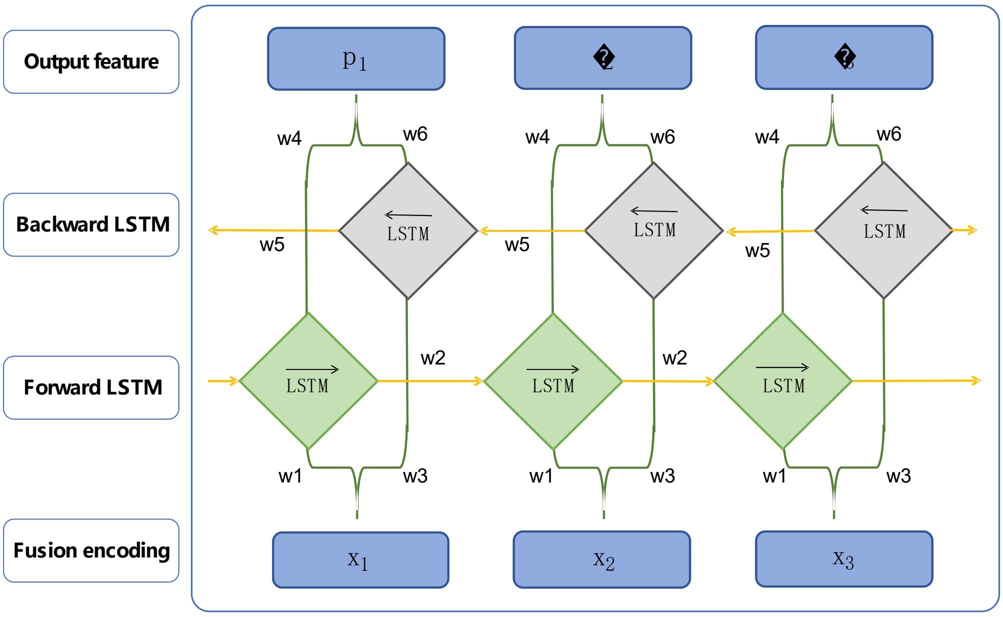


**Supplementary Figure 1. Bi-LSTM Architecture.** It can be shown that the forward layer and the backward layer are connected to the output layer, which contains shared weights w1-w6. In the forward layer, the forward calculation is performed from time 1 to time t, and the output of the forward hidden layer at each time is obtained and saved. In the backward layer, reverse the calculation from time t to time 1 to get and keep the output of the backward hidden layer at each time. Finally, the final output is obtained by combining the output results of the corresponding forward layer and backward layer at each time.

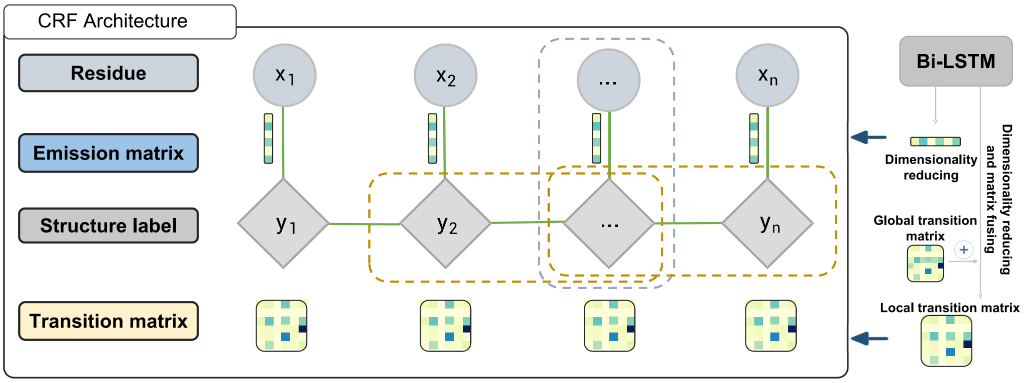


**Supplementary Figure 2. CRF Architecture**. The emission matrix consisting of the possibility of different sub-structures at each residue can be learned by Bi-LSTM layer. The local transition matrix is the fusion of the global transition matrix and the residue features from Bi-LSTM for transformation scoring among sub-structures.

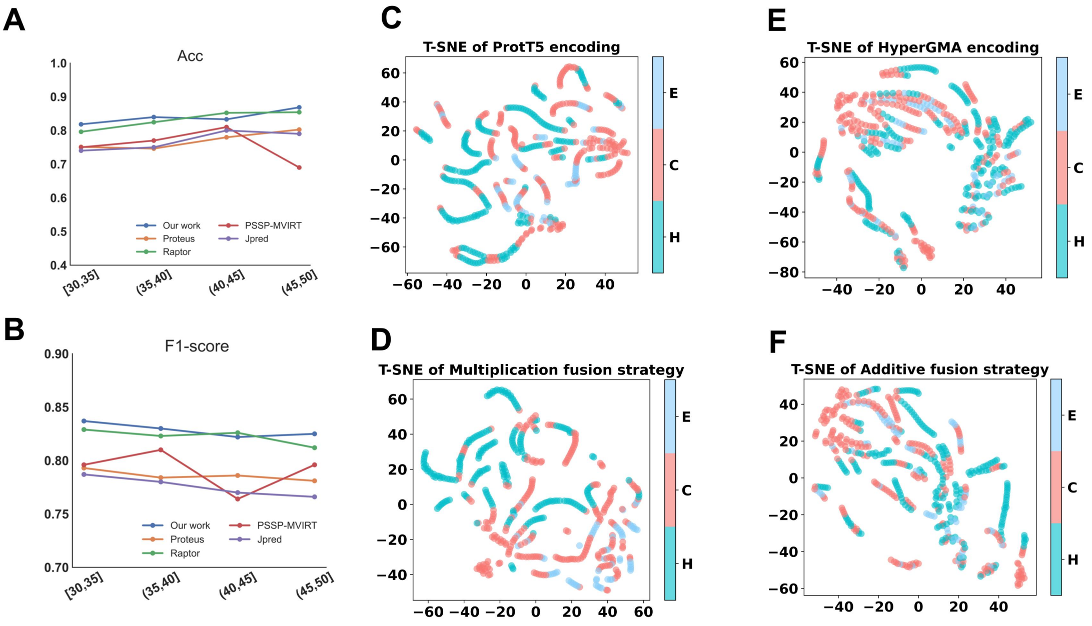


**Supplementary Figure 3.** (**A**) Acc is used as the evaluation metric; (**B**) F1-score is used as the evaluation metric. (**C–F**) represent t-SNE visualization results of the fused extractors in multiplication or additive and individual extractors of ProtT5, HyperGMA, respectively.

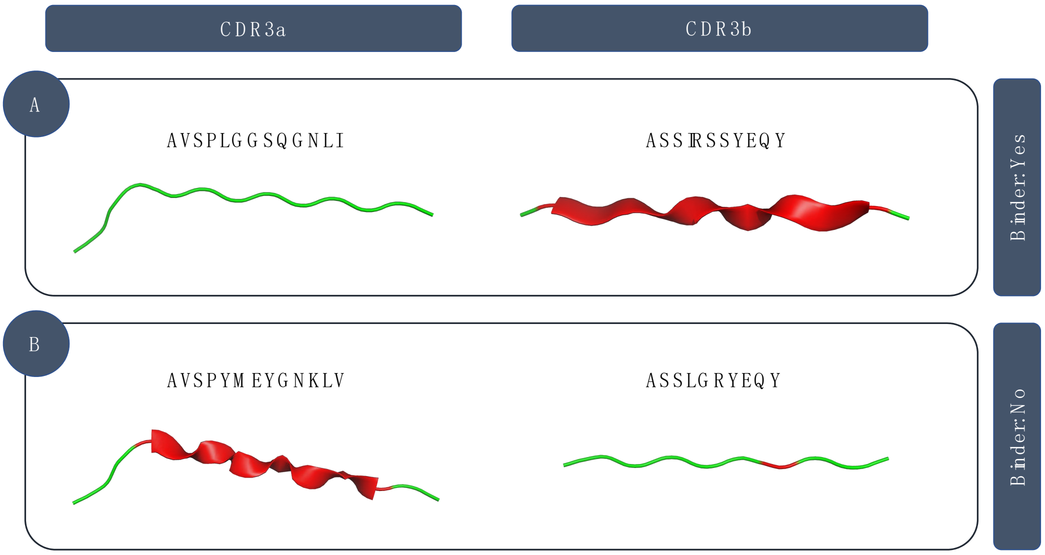


**Supplementary Figure 4.** Visualization of the secondary structures of the two peptide sequences predicted by our method.

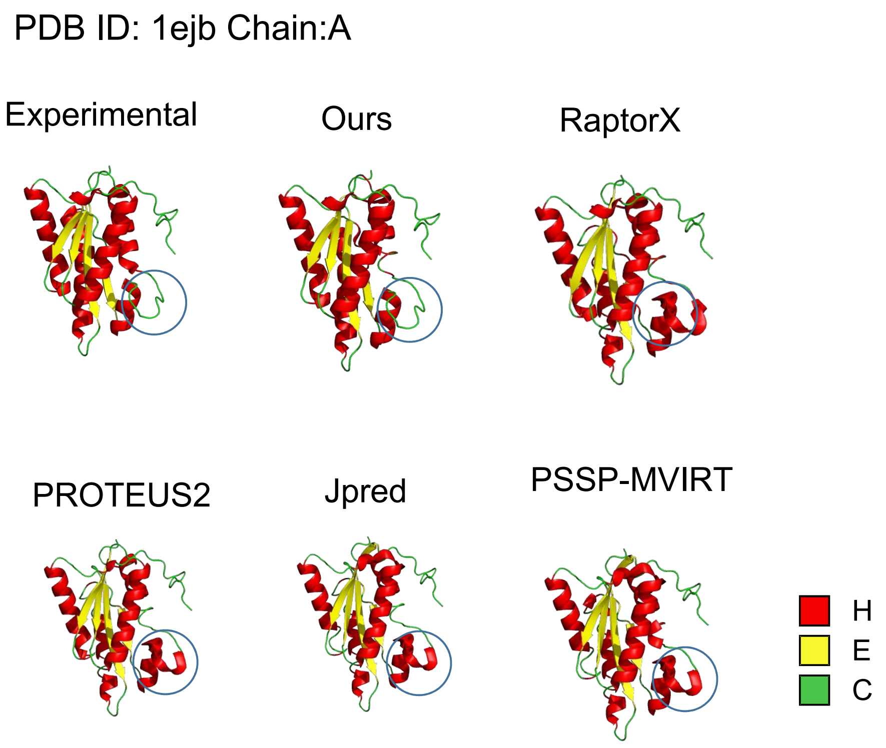


**Supplementary Figure 5. Visualization of secondary structures mapped into tertiary structures for our method and existing methods including RaptorX, PSSP-MVIRT, PROTEUS2 and Jpred.** The visualization of predictions by our method and existing methods for the peptide with PDB ID: 1ejb. The prediction regions with large differences from different methods are marked with circles.

### Experimental settings

To train a robust and accurate model, we apply the layer normalization and dropout techniques. The layer normalization is used in integrating the features from the pretrained model ProtT5 and HyperGMA. Layer normalization can impose constraints on the "scale" problem, which may be caused by the embedding of multiple features in the learning process, effectively reducing the model variance. As for dropout, it is inserted into the attention layer of HyperGMA, solving the overfitting problem.

During slicing peptides into fragments and dividing fragments into residue groups to construct the structure of the hypergraph, we set 12 residues long as the length of the fragment, and there are four residues coincident between two neighboring fragments. As for the residues group, two residues are used to form a group with one same residue in the neighboring two groups. In our study, the whole deep learning models were trained globally by the Adam algorithm with a learning rate l = 1e-4 to minimize the cost function Loss. The training epoch is set to 200, and it performs best in the around 121 epoch. All the training and testing procedures were performed based on Nvidia RTX 3090 GPUs and implemented by python based on PyTorch.

### Supplementary metrics

To evaluate the results of comparison in prediction of peptide toxicity, we used six traditional evaluation metrics commonly used in binary classification tasks, including Sensitivity (SN), Specificity (SP), False discovery rate (FDR), False positive rate (FPR), Accuracy (ACC) and Mathew's correlation coefficient (MCC). The metrics are calculated as follows:

$$SN = \frac{TP}{TP + FN} \tag{1}$$

$$SP = \frac{TN}{TN + FP} \tag{2}$$

$$FDR = \frac{FP}{TP + FP} \tag{3}$$

$$FPR = \frac{FP}{FP + TN} \tag{4}$$

$$ACC = \frac{TP + TN}{TP + TN + FP + FN} \tag{5}$$

$$MCC = \frac{TP \times TN - FP \times FN}{\sqrt{(TP + FN)(TP + FP)(TN + FP)(TN + FN)}} \tag{6}$$

where TP (true positive) and TN (true negative) represent the numbers of correctly predicted positive samples and negative samples, respectively; FP (false positive) and FN (false negative) represent the numbers of incorrectly predicted positive samples and negative samples, respectively. The metric SN measures the prediction ability of a predictor for positive samples, while the metric SP measures the ability of the predictor for negative samples. FDR calculates the proportion of errors in the positive samples predicted by the predictor, while FPR calculates the proportion of negative samples that are mistaken as positives by the predictor. ACC and MCC are used to evaluate the overall performance of a predictor. Moreover, the ROC (receiver operating characteristic) curve and PR (precision-recall) curve are often used to intuitively evaluate the overall predictive performance of a predictor. Here, we calculated the area under the ROC curve (AUC) to assess the overall predictive performance. The value of AUC is from 0.5 to 1. The larger the value of AUC, the better and more robust performance.
